# The herpetofauna of Despotiko Island (Cyclades, Greece)

**DOI:** 10.1101/2025.03.06.641418

**Authors:** Jennifer Rose Poole, Joshua Smith, Thomas Hesselberg, Panayiotis Pafilis

**Affiliations:** Department of Biosciences, University of Exeter, Exeter, Devon, EX4 4RN, UK; Jersey Internation Centre of Advanced Studies, St. Helier, Jersey, Channel Islands; Department of Biology, University of Oxford, Oxford, UK; Coiba Scientific Station, Panama City, Panama; Zoological Museum, National and Kapodistrian University of Athens; Section of Zoology and Marine Biology, Department of Biology, National and Kapodistrian University of Athens, Athens, Greece

**Keywords:** reptiles, Aegean, islands, archaeological sites, *Lacerta citrovittata*, *Eryx jaculus*, Greece

## Abstract

The Aegean islands are a known hotspot for herpetofauna and have been extensively studied in this area. However, there are still numerous islands that lack this research. This includes the uninhabited islet of Despotiko found in the Paros archipelago, Cyclades. It is known for its importance in archaeology, which in turn attracts tourism. However, there is not yet a complete understanding on the presence of herpetofauna there, with the last published records written in 1977. In an effort to address this, a combination of past *ad-hoc* sightings and visual surveys were carried out during summer 2024. These yielded two new records of reptile species, *Eryx jaculus* and *Lacerta citrovittata*. Furthermore, the enclosed archaeological site had greater reptile diversity and richness than outside the site. This highlights the potential importance of the archaeological site as a key reptile habitat. Our findings help to improve our understanding of reptilian diversity in the archipelago, providing avenues for further research into the potential interactions between archaeological sites and reptile diversity.

## Introduction

The Aegean islands in Greece are famous for hosting an impressive herpetofauna that includes numerous endemic species (Werner 1930; Lymberakis et al. 2018). Thanks to their unique biodiversity, early naturalists as well as contemporary researchers show a strong preference for these islands (e.g. Bedriaga 1883; Broggi 2024). Therefore, the Aegean islands, the “showcase” of the Greek herpetofauna (Christopoulos et al. 2019) are well studied and the herpetofauna of even tiny islets has been documented (e.g. Buchholz 1962; Pafilis et al. 2024a). Interestingly, however, a few islands have somehow eluded this trend of herpetological meticulousness. Despotiko (Greek: *Δεσπ0τIκό*, deriving from *δεσπότησ* meaning lord, ruler), an island in the Cyclades group, is such a case.

### Study Area

Despotiko is a small island located in the central Aegean Sea (36° 57’ 45.3126’’ N, 25° 0’ 12.1998’’ E, WGS84 datum; Fig. 1). It encompasses an area of ∼7.6 km^2^ with high topographical variation and a peak elevation of ∼187 m above sea level. Metamorphic rocks (gneiss and marble) cover most of the island (Anastopoulos 1963). Surrounding islands include Antiparos, Tsimintiri and Strongyli (Fig. 1), which are separated by shallow waters ranging from 1 – 4 m in depth. Paleogeographic reconstruction revealed that Despotiko used to be connected to neighbouring islands, forming a mega-island with Strongyli, Antiparos, Tsimintiri, Paros and Naxos ca. 8,000 B.P. (Kapsimalis et al. 2009). Subsequent sea level rise over time has led to the separation of islands to the present-day configuration. The climate surrounding Despotiko is a typical Mediterranean climate, with hot, dry summers and cooler, wetter winters (Archibold 1995). There are, however, Meltemi/Etesian winds which are strong, dry, north winds that blow mid-May to mid-September. These can make the sea rough which can restrict access to the island and limit reptile activity (Buttle 1993).

**Figure 1.**
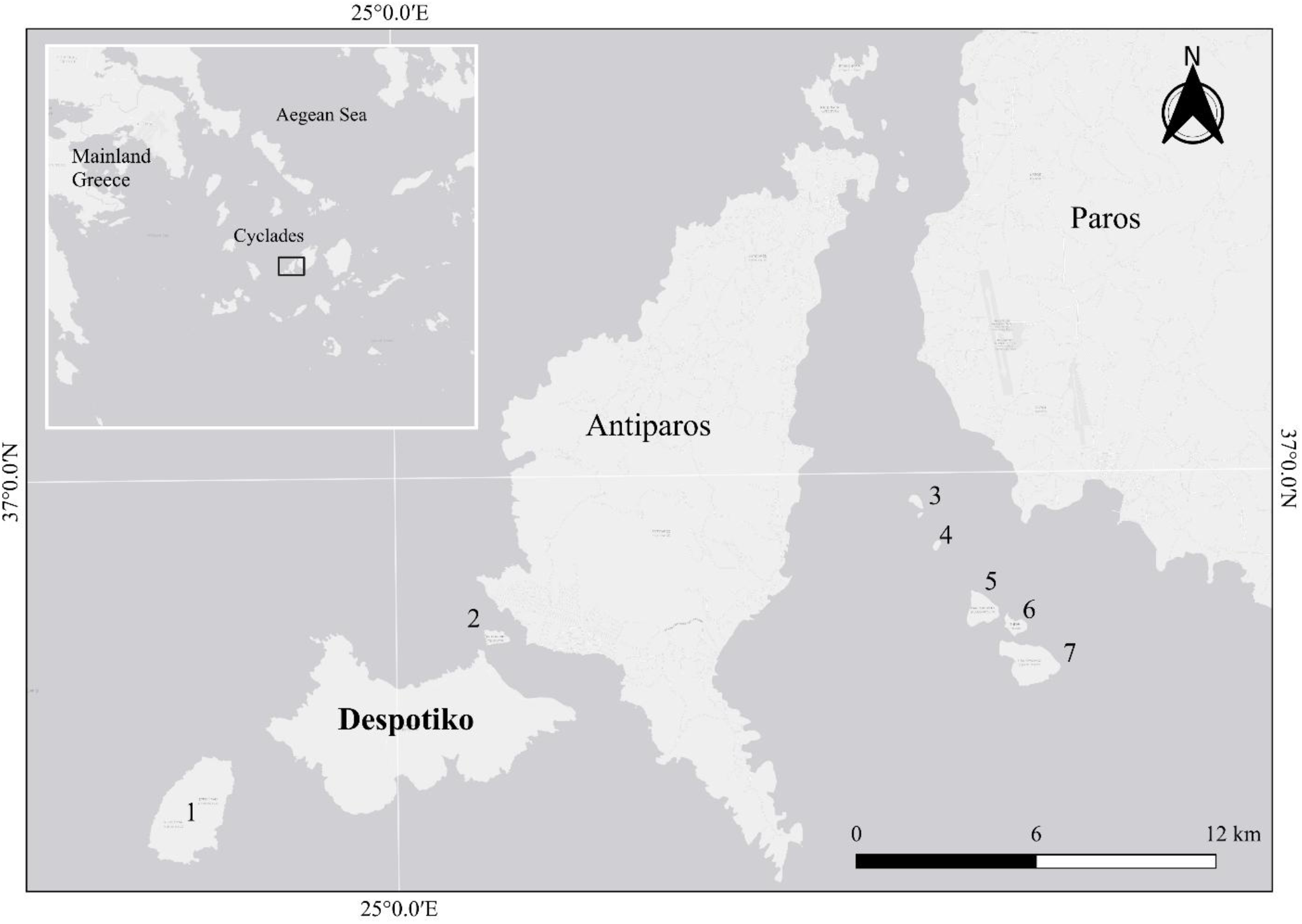
Location of Despotiko in relation to other islands in the Cyclades: 1 – Strongyli, 2 – Tsimintiri, 3 – Tourna, 4 – Preza, 5 – Glaropounta, 6 – Tigani, 7 – Panderonisi.

The island is uninhabited today, but remains of pottery and structure foundations suggest the island was extensively inhabited in the past (Broodbank 2000). Indeed, people settled Prepesinthos (the ancient name for Despotiko) from the early Bronze Age and it seems that human presence was continuous till late 17^th^ century (Sphyroeras et al. 1985). The Sanctuary of Apollo was a religious regional centre in the Cyclades and flourished during the Archaic period (600–500 BCE). This may have influenced the flora and fauna on the island.

Nowadays, the main human activities are archaeological excavations and goat farming (∼200). Goat grazing is widespread, heavy and continuous across the island. The exception is the archaeological site which is situated on the north-east side of the island, surrounded by fences to prevent unauthorised access and goat grazing. Despotiko also attracts daily cruise tourism, which is a growing industry within the area. The island has various ravines, pockets of dense vegetation and hilly areas. The goats are within an enclosed area, but the stockmen often release them, allowing them free rein across the island. The vegetation consists of typical Mediterranean phrygana, dominated by *Pistacia lentiscus* and *Juniperus turbinata*, with clusters of *Erica manipuliflora* on rocky, dry soil (Fig. 2). The entire island is covered by the Natura 2000 network with two sites (GR4220025 and GR4220017).

**Figure 2.**
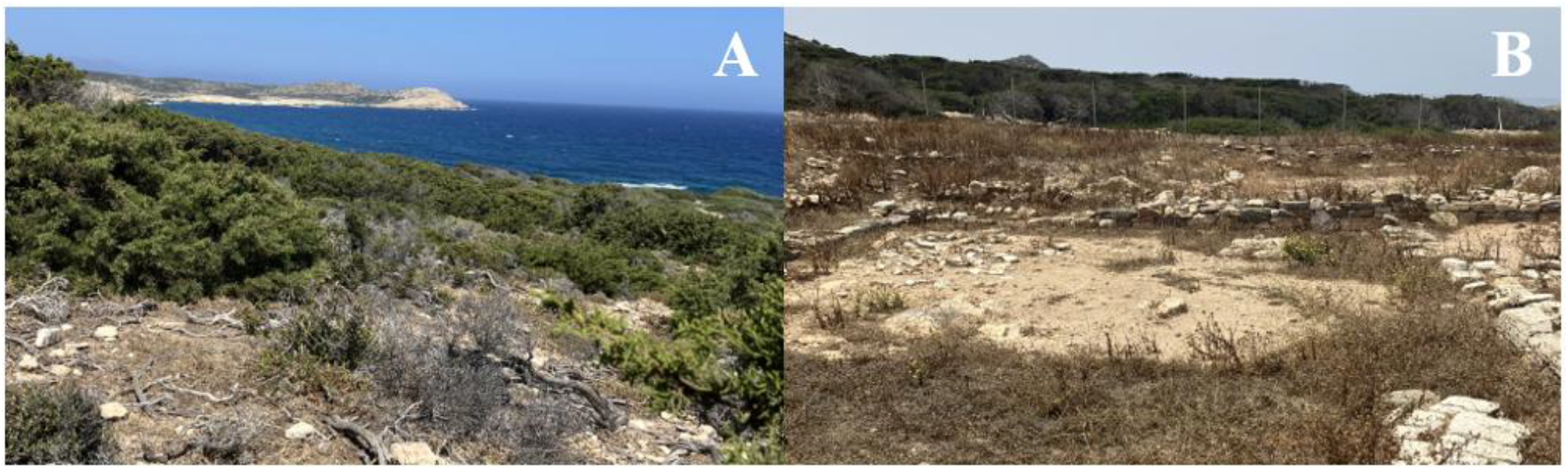
Photo of vegetation found on Despotiko outside the archaeological site (A) and inside the archaeological site (B).

Records of fauna on Despotiko are relatively limited. However, a survey carried out by Gruber and Fuchs (1977) across the Paros archipelago determined 13 species of herpetofauna, of which five were found in Despotiko. These included *Mediodactylus kotschyi* (Kotschy’s gecko), *Hemidactylus turcicus* (Mediterranean house gecko), *Ablepharus kitaibelii* (European copper skink), *Laudakia stellio* (Agama lizard) and *Natrix natrix* (grass snake). These findings were verified by a more recent study (Itescu et al. 2019). Neither amphibians, nor the reptiles *Podarcis erhardii* (Aegean wall lizard), *Elaphe quatuorlineata* (four-lined snake), *Telescopus fallax* (European cat snake), *Vipera ammodytes* (nose-horned viper) and *Eryx jaculus* (javelin sand boa) were found on Despotiko despite being present elsewhere in the Paros archipelago (Pafilis and Maragou 2020). Bird life is scarce, while the main mammals on the island, besides goats, are introduced rats which may predate on lizard<u>s</u> and their eggs (Schwarz et al. 2020).

## Methods

The island was surveyed between 17/06/2024 – 23/06/2024. A special permit was issued by the Ministry of Environment and Energy (AΔA: 6ZΣΘ4653Π8-ΨO9) to survey outside the archaeological site, and by the Ministry of Culture to survey within the archaeological site. Visitation to the island was restricted by boat access and was only permitted when archaeologists were present. Surveying was split across two 200 × 200 m areas covering the archaeological site and just outside (Fig. 3). Reptiles were surveyed by 35- minute walking transects and flipping rocks. 10 coordinates at least 50 m apart were randomly generated using QGIS and these were used to determine the transect route within the area. The first author was the main observer throughout the survey to reduce bias. Five rocks greater than 15 cm diameter were randomly selected and flipped every five minutes along the transect. Due to visitation/time restrictions based on boat times, only two surveys per area were conducted (one morning and one afternoon on different days to account for variability in reptile activity) totalling four surveys carried out on the island. Only the north- eastern side of the island was surveyed which was closest to the goat farm and has the greatest grazing pressure. Therefore, reptiles not found in this study may be present in other habitat types on the island. Visual sightings were recorded using ArcGIS Survey123 (accuracy ± 4m). Temperature and wind speed were recorded at eye level using a HoldPeak anemometer at the start and end of each survey. Reptiles were identified visually using existing photographs and where possible, photographs were taken. No reptiles were captured or handled during the study.

**Figure 3.**
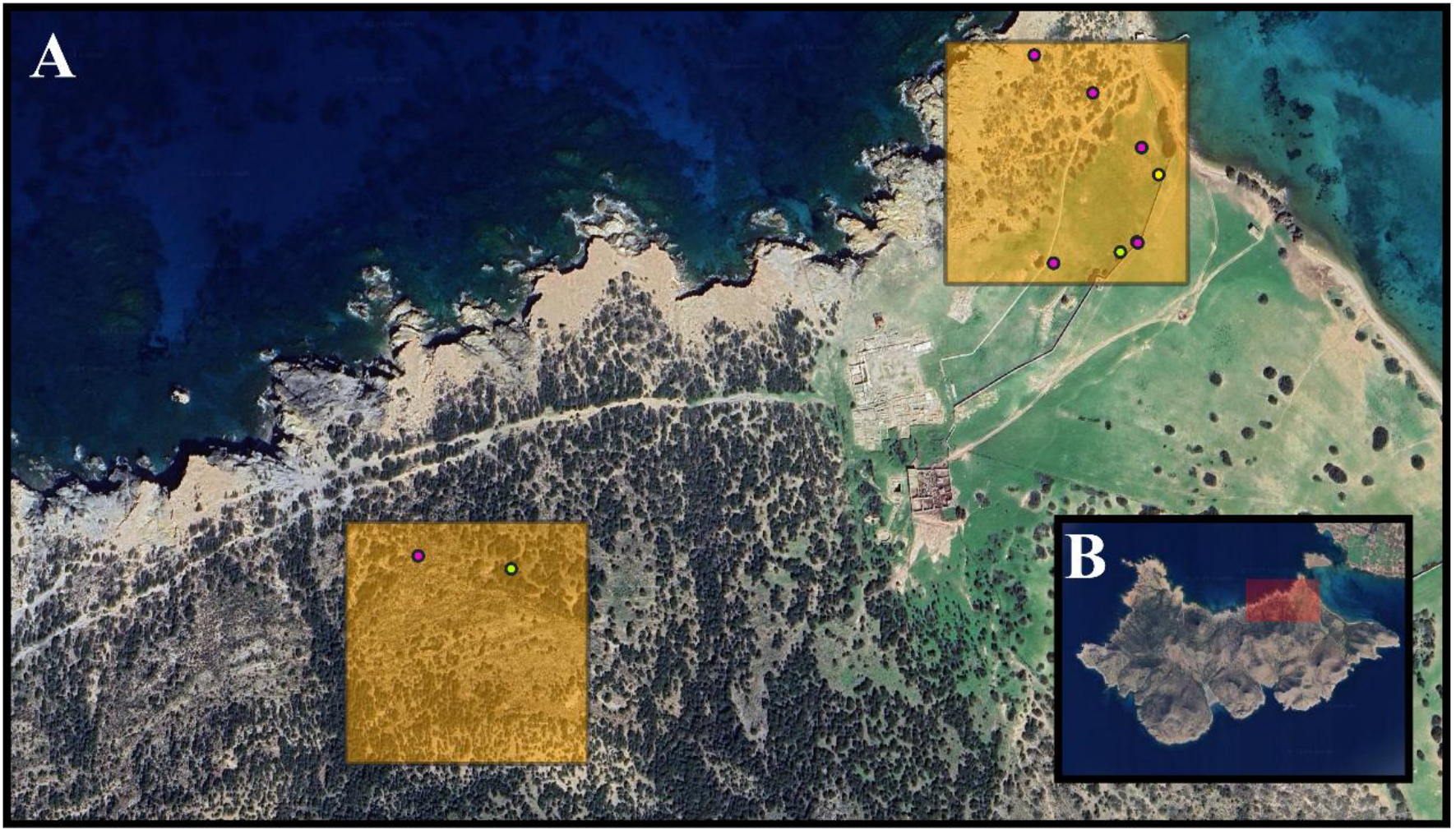
A) Survey sites on Despotiko with sightings of *Mediodactylus kotschyi* (purple), *Lacerta citrovittata* (green) and *Ablepharus kitaibelii* (yellow); B) Area of Despotiko the survey sites were located.

Reptile diversity and richness were calculated from finding the average Shannon’s diversity and total species richness from morning and afternoon surveys across the two sites using the Vegan package in R Studio (Oksanen et al. 2022; R Core Team 2023). An insufficient number of observations were made to conduct statistical analyses however this could be explored in a future study over a longer survey period or a larger dataset.

## Results

The herpetofauna found on Despotiko (previous and observed records from this study) consists of the following species:

### *Mediodactylus kotschyi* (Steindacher 1870) (Fig. 4A)

Despotiko - first published record by Gruber and Fuchs (1977); other published records by Itescu et al. (2017) and Schwarz et al. (2020); records in this study: 10 sightings all found under rocks. This was the most common reptile found on the island.

**Figure 4.**
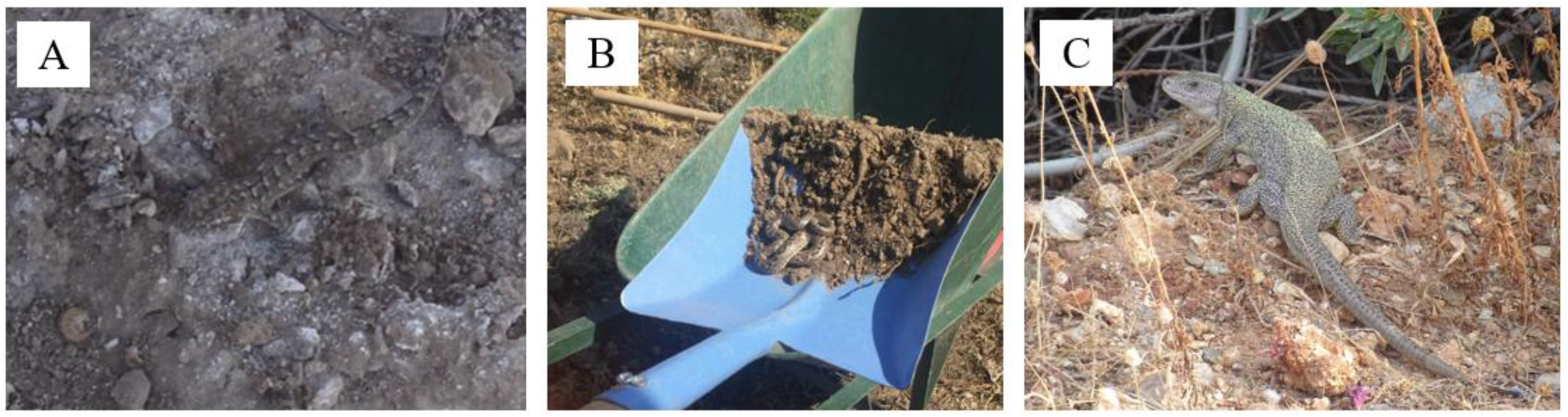
A) *Mediodactylus kotschyi*; B) *Eryx jaculus* (Credit: Wilkie, 2024, pers. comm.); C) *Lacerta citrovittata*.

### *Hemidactylus turcicus* (Linnaeus 1758)

Despotiko – first published record by Gruber and Fuchs (1977); other published records by Itescu et al. (2017); records in this study – none. This species is common around human settlements and although it was not observed in this study, this may be due to it being present on the archaeological structures themselves which were not included in the transect.

### *Ablepharus kitaibelli* (Bibron & Bory de Saint-Vincent 1833)

Despotiko – first published record by Gruber and Fuchs (1977); records in this study - one individual found along transect in the archaeological site under a rock, one ad-hoc sighting found outside the archaeological site next to *E. manipuliflora*.

### *Laudakia stellio* (Linnaeus 1758)

Despotiko – first published record by Gruber and Fuchs (1977); records in this study – none. Only one specimen was recorded by Gruber and Fuchs (1977), highlighting the rarity of the species. *Laudakia stellio* prefers dry habitats with stone walls and rocks to bask in the sun. The lack of *L. stellio* from Despotiko is surprising as it is a common, and conspicuous, species in the neighbouring Antiparos, Paros and Naxos.

### *Natrix natrix* (Linnaeus, 1758)

Despotiko – first published record by Gruber and Fuchs (1977); records in this study – none. *Natrix natrix* prefer more aquatic, freshwater habitats with amphibians being a key food source. The lack of water and presence of amphibians may lead to this island being unsuitable for maintaining *N. natrix* populations. The specimen reported by Gruber and Fuchs (1977) could be an accidental finding as grass snakes are known to cover short distances even at sea: few years ago, Mossman et al. (2016) recorded a melanistic *N. natrix* on the nearby small islet Tigani, between Paros and Antiparos.

### *Eryx jaculus* (Linnaeus, 1758)

Despotiko – first published record; other records – an *ad-hoc* sighting by a group of archaeologists. Found curled under a white rock at 9:18am on 11/06/2019 (Wilkie, 2024, pers. comm; Fig. 4B); records in this study – none. The javelin sand boa follows a cryptic behavioural pattern and thus is rarely observed. Reptiles in Greece often use archaeological sites as hibernacula and when excavations resume in the spring it is quite common for archaeologists to encounter snakes (Pafilis et al. 2024b).

### *Lacerta citrovitatta* (Werner 1938) (Fig. 4C)

Despotiko – first published record; other records – none; records in this study – two individuals found during the reptile surveys (one per area) plus two ad-hoc sightings within the archaeological site.

## Discussion

Overall species diversity and richness were higher in the archaeological site compared to outside this area (Table 1). This may highlight the benefit of archaeological sites as a key habitat for reptiles on the island, and the possible detrimental effects of goats that wander outside the site. This has the potential to be explored further and more in-depth. It has been evidenced that goat presence on islands in the Skyros archipelago had a greater negative impact on the endemic Skyros lizard (*Podarcis gaigae*) compared to islands with no goats (Pafilis et al. 2013). Comparisons between Despotiko and the neighbouring island Strongyli, which is goat-free, could shed more light on the impacts of goat grazing. Additionally, archaeological sites have been shown to have high plant species richness in the Peloponnese (Panitsa et al. 2024), which may contribute to higher richness in other taxa such as reptiles.

**Table 1.** A comparison of habitat, Shannon’s diversity and richness across the two survey sites.

| Survey site | Habitat | Mean Shannon's diversity ($H'$) | Total species richness |
| --- | --- | --- | --- |
| Non-archaeological | Dry, dense pockets of *Juniperus turbinata*, *Pistacia lentiscus* and *Erica manipuliflora*. | 0 | 1 |
| Archaeological | Dry, sparse vegetation consisting of *Hirschfelda incana*, *Cichorium sp.* and *Carlina corymbosa*. Scattered marble rocks. | 0.399 | 2 |

Plant species richness may be impacted in Despotiko as despite the fencing, goats occasionally enter the archaeological site. Higher fencing around the area could mitigate this. Despotiko is also much smaller than many other islands in the Aegean and may follow the pattern in the island species-area relationship where larger islands tend to have greater species richness due to having a greater diversity of habitats which increases niche availability and therefore can support more species. Similarly, this pattern was also seen in the Sporades where islands of similar size to Despotiko had similar reptile species richness as found in this study (Foufopoulos et al. 2024). Likewise, Folegandros was also noted to have lower reptile richness than expected, possibly due to the barren, arid landscape (Itescu et al., 2017). This reason may also contribute to the low species richness found on Despotiko.

## Conclusion

150 years of herpetological research yielded a detailed description of the Aegean herpetofauna (Pafilis 2010; Lymberakis et al. 2018), which is not, however, exhaustive. New records from the islands further enhance a growing body of literature, either with reports for new species (e.g. Grano and Cattaneo 2020; Pafilis and Kapsalas 2024) or for small islets’ herpetofaunas (e.g. Tzoras et al. 2019; Pafilis et al. 2020). In this context, our study on Despotiko herpetofauna, with two new records, comes to improve our knowledge and understanding of reptilian diversity of the archipelago.

## Acknowledgements

The authors would like to thank the Greek Ministry of Environment and Energy and Yannos Kourayos from the Ministry of Culture for permission and access to survey Despotiko.

Special thanks to Caitlin Horne for her assistance in the field. Jersey International Centre of Advanced Studies (University of Exeter) partially funded the study.

## Notes

### Competing Interest Statement

The authors have declared no competing interest.

